# The receptor-like kinases BAM1 and BAM2 promote the cell-to-cell movement of miRNA in the root stele to regulate xylem patterning

**DOI:** 10.1101/603415

**Authors:** Pengfei Fan, Hua Wang, Hao Xue, Tabata Rosas-Diaz, Weihua Tang, Heng Zhang, Lin Xu, Rosa Lozano-Duran

## Abstract

Xylem patterning in the root is established through the creation of opposing gradients of miRNAs and their targets, enabled by the cell-to-cell spread of the former. The miRNAs involved in xylem patterning, miR165/6, move through plasmodesmata, but how their trafficking is regulated remains elusive. Here, we describe that the receptor-like kinases BAM1/2 are required for the intercellular movement of miR165/6 in the stele and hence proper xylem patterning in the root.

## MAIN TEXT

Tissue patterning in plant organ development depends primarily on positional information, which must be communicated between cells. Different mobile molecules can mediate cell-to-cell communication, including phytohormones, transcription factors, or peptides. In the past decade, multiple works have uncovered the relevance of small non-coding RNAs (sRNA) as mobile signaling molecules capable of acting as morphogens in plant development, determining leaf polarity, root vascular patterning, embryo meristem formation, female gametogenesis, and maintenance of the shoot apical meristem, regulating the acquisition of cell fate in a dose-dependent fashion (reviewed in ^1-3^). Interestingly, it has been recently shown that the cell-to-cell movement of microRNAs (miRNAs) is directional ^4^, indicating that this process must be precisely regulated.

An elegant example of how sRNA can determine pattern formation is provided by the study of xylem patterning in the root. Xylem patterning is established by a robust regulatory pathway comprising bidirectional cell signaling mediated by miRNAs 165 and 166 (miR165/6) and the transcription factors SHORT ROOT (SHR) and SCARECROW (SCR) ^5^: xylem precursors differentiate into two types of xylem vessels: metaxylem cells, with pitted secondary cell walls, in the centre of the vascular cylinder, and protoxylem cells, distinguishable by their spiral walls, in a peripheral position (Figure 1A). SHR is produced in the steel, and moves from cell to cell to the endodermis, where it activates *SCR* and, together with the latter, *MIR165a* and *MIR166b*. The resulting miR165 and miR166 move into the stele to pattern the *class III HOMEODOMEIN-LEUCIN ZIPPER* (*HD-ZIP III*) mRNA domains, particularly that of *PHABULOSA* (*PHB*), restricting them to the centre of the stele, which results in correct xylem patterning with formation of both metaxylem and protoxylem ^5,6^ (Figure 1A). Although miR165/6 have been shown to move symplastically through plasmodesmata^7^, how their trafficking is regulated remains elusive.

**Figure 1.**
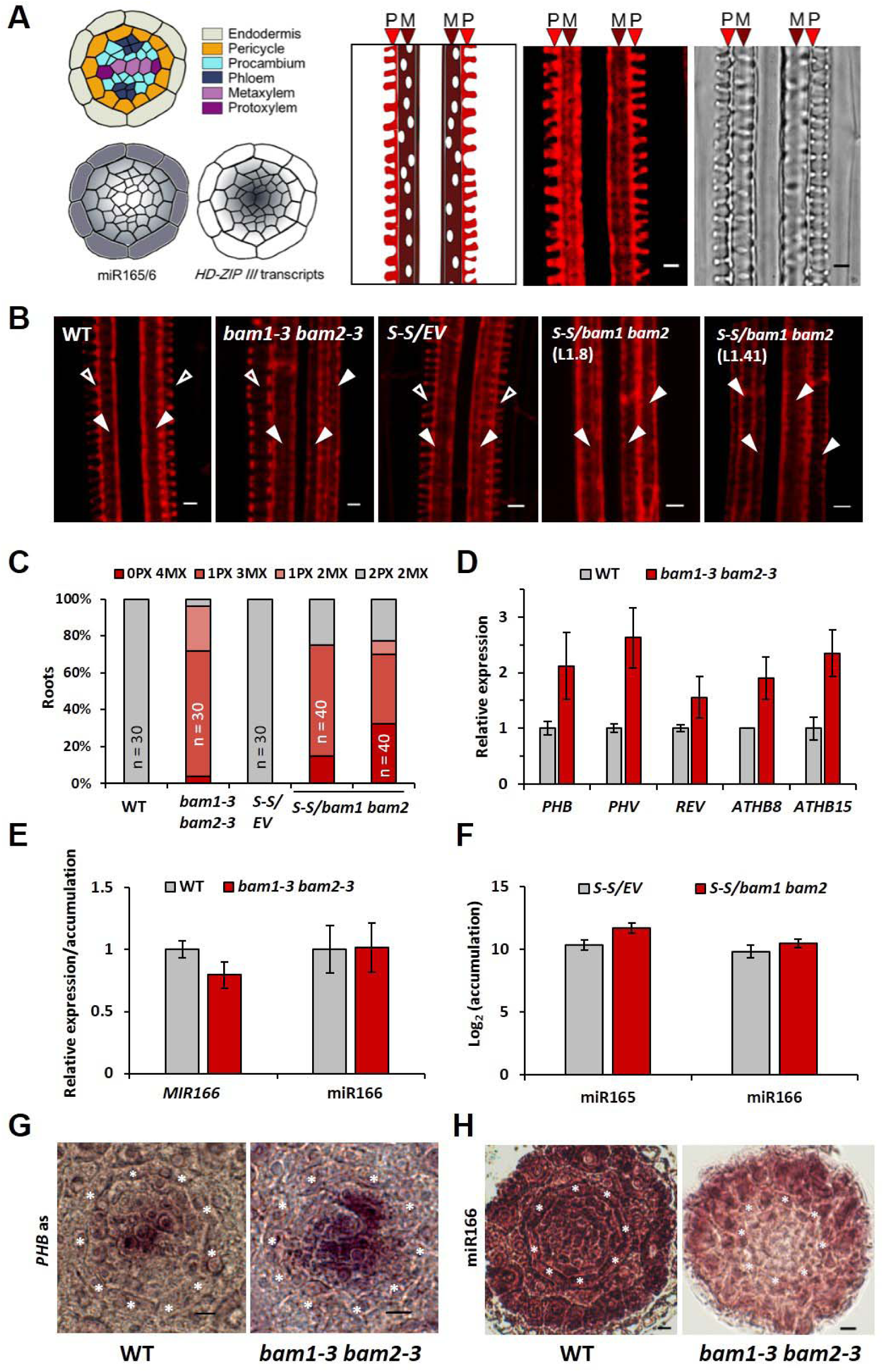
BAM1 and BAM2 play a redundant role in xylem development through the promotion of the cell-to-cell movement of miR165/6. **A.** Schematic representations of a cross-section of the Arabidopsis root stele surrounded by the endodermis, distribution of miR165/6 and *HD-ZIP III* transcripts, and typical structure of protoxylem (P) and metaxylem (M) in a longitudinal section of the root; on the right, confocal image of a longitudinal section of the root showing basic fuchsin-stained protoxylem (P) and metaxylem (M), and bright field image of the same region. Scale bar = 3µm. **B.** Confocal micrographs of basic fuchsin-stained xylem in control and *bam1 bam2* six-day-old roots. Empty arrowheads indicate protoxylem; filled arrowheads indicate metaxylem. WT: wild type (L*er*); S-S: *SUC:SUL*; EV: empty vector. Scale bar = 3µm. **C.** Quantification of the number of protoxylem (PX) and metaxylem files (MX) in the roots of listed genotypes. **D**, **E.** Accumulation of transcripts of the *HD-ZIPIII* family genes (**D**), transcripts of *MIR166*, and miR166 (**E**) in WT and *bam1-3 bam2-3* double mutant six-day-old seedlings as measured by qRT-PCR. Results are the average of three biological replicates. Error bars indicate SD. **F.** miR165/6 accumulation in *S-S/bam1 bam2* double mutants and SUC-SUL/EV control as measured by sRNA-seq. **G.** *In situ* hybridization with a *PHB* mRNA specific probe on cross-sections of WT and *bam1-3 bam2-3* roots. Seven roots were checked per genotype; all roots showed a similar phenotype. One representative picture is shown. **H.** *In situ* hybridization with a miRNA166-specific LNA probe on cross-sections of WT, *bam1-3 bam2-3* roots. Ten roots were checked per genotype; five *bam1-3 bam2-3* roots showed the miR166 distribution pattern displayed in this figure, with weaker signal in the stele. Scale bar = 6µm. Asterisks indicate the position of the endodermis.

The plasma membrane- and plasmodesmata-localized receptor-like kinases BARELY ANY MERISTEM (BAM) 1 and 2 have been recently described as required for the cell-to-cell spread of RNA interference (RNAi) in the reporter *SUC-SUL* plants ^8^, in which production of mobile siRNA against the endogenous *SULPHUR* (*SUL*) gene in phloem companion cells causes non-cell autonomous silencing observable as a chlorotic phenotype around the leaf veins ^9^. Whether BAM1/2 also play a role in the cell-to-cell movement of other sRNAs, such as miRNAs, is yet to be determined.

In Arabidopsis roots, *BAM1* is strongly and specifically expressed in the stele (Figure S1). We hypothesized that, considering this particular expression pattern, if BAM1 regulates movement of miRNA, it could mediate the cell-to-cell spread of miR165/6, hence acting as a regulator of xylem patterning. In order to determine whether BAM1/2 are required for correct xylem formation in the root, we observed xylem patterning in *bam1/2* mutants ^9,10^. Interestingly, *bam1 bam2* double mutants, but not *bam1* or *bam2* single mutants, display shorter roots (Figure S2) and show xylem defects consistent with a malfunction of miR165/6, namely absence of protoxylem files and overproliferation of metaxylem at the expense of protoxylem (Figure 1B and C; Figure S3). At the molecular level, *bam1 bam2* mutants display increased levels of *HD-ZIP III* transcripts, but are not affected in the expression of *MIR165/166, SHR*, or *SCR*, or in the accumulation of miR165/6 (Figure 1D-F; Figure S4). Further supporting the idea that movement of miR165/6 is affected in the double mutants, the distribution of the *PHB* transcript is less restricted in the stele in the absence of BAM1/2 (Figure 1G), while lower levels of miR166 can be detected in this area (Figure 1H). On the contrary, transgenic plants overexpressing *BAM1* have normal xylem and wild type-like accumulation of the *HD-ZIP III* transcripts (Figure S5). Taken together, these results indicate that BAM1/2 are required for proper xylem patterning, likely due to a function as positive regulators of the cell-to-cell movement of miR165/6.

The C4 protein from the geminivirus *Tomato yellow leaf curl virus* (TYLCV) interacts with the intracellular domain of BAM1/2 at the plasma membrane and has a negative impact on the cell-to-cell spread of RNAi ^9^. In order to see whether the activity of C4 can have an effect of xylem patterning, we observed the xylem in roots of transgenic plants expressing C4 under the control of the constitutive 35S promoter ^9^. Strikingly, expression of C4 led to xylem defects similar to those observed in *bam1 bam2* mutants (Figure 2A and B). Plasma membrane localization of C4 is essential for this phenotype, since plants expressing the mutated version C4_G2A_, which loses its membrane association and localizes to chloroplasts exclusively ^9^, have wild type-like xylem (Figure S6). Transgenic plants expressing C4, but not C4_G2A_, have increased levels of *HD-ZIP III* transcripts (Figure 2C, Figure S6). However, C4 does not affect the expression of *MIR166, SHR*, or *SCR*, or the accumulation of miR165/6 (Figure 2D and E; Figure S7). As observed for *bam1 bam2* mutants, the distribution of the *PHB* transcript in the stele is broader in the C4-expressing plants (Figure 2F), and lower levels of miR166 are detected in this part of the root (Figure 2F).

Since miR165/6 are produced in the endodermis, and from here traffic inwards into the stele establishing a gradient that determines *HD-ZIP III* dosage ^5^, we reasoned that if C4 is exerting its effect on xylem patterning through the interference with the cell-to-cell movement of miRNAs, then expressing C4 in the endodermis layer should have a non-cell autonomous effect and be sufficient to cause the observed phenotype. Indeed, transgenic plants expressing C4 under the control of the endodermis-specific *SCR* promoter display xylem patterning and related molecular phenotypes similar to those previously described for *bam1 bam2* and *35S:C4* transgenic lines (Figure 2A-D; Figure S7), including a wider *PHB* domain and lower miR166 in the root stele (Figure 2F, G). Moreover, despite its localization in plasmodesmata ^9^, C4 does not disturb the movement of SHR-GFP (Figure S8). Therefore, C4 interferes with xylem patterning non-cell-autonomously, most likely through an impairment of miR165/6 movement. Of note, transgenic plants expressing C4 under the SCR promoter display wild type-like rosettes, but abnormal floral stems (Figure S9).

**Figure 2.**
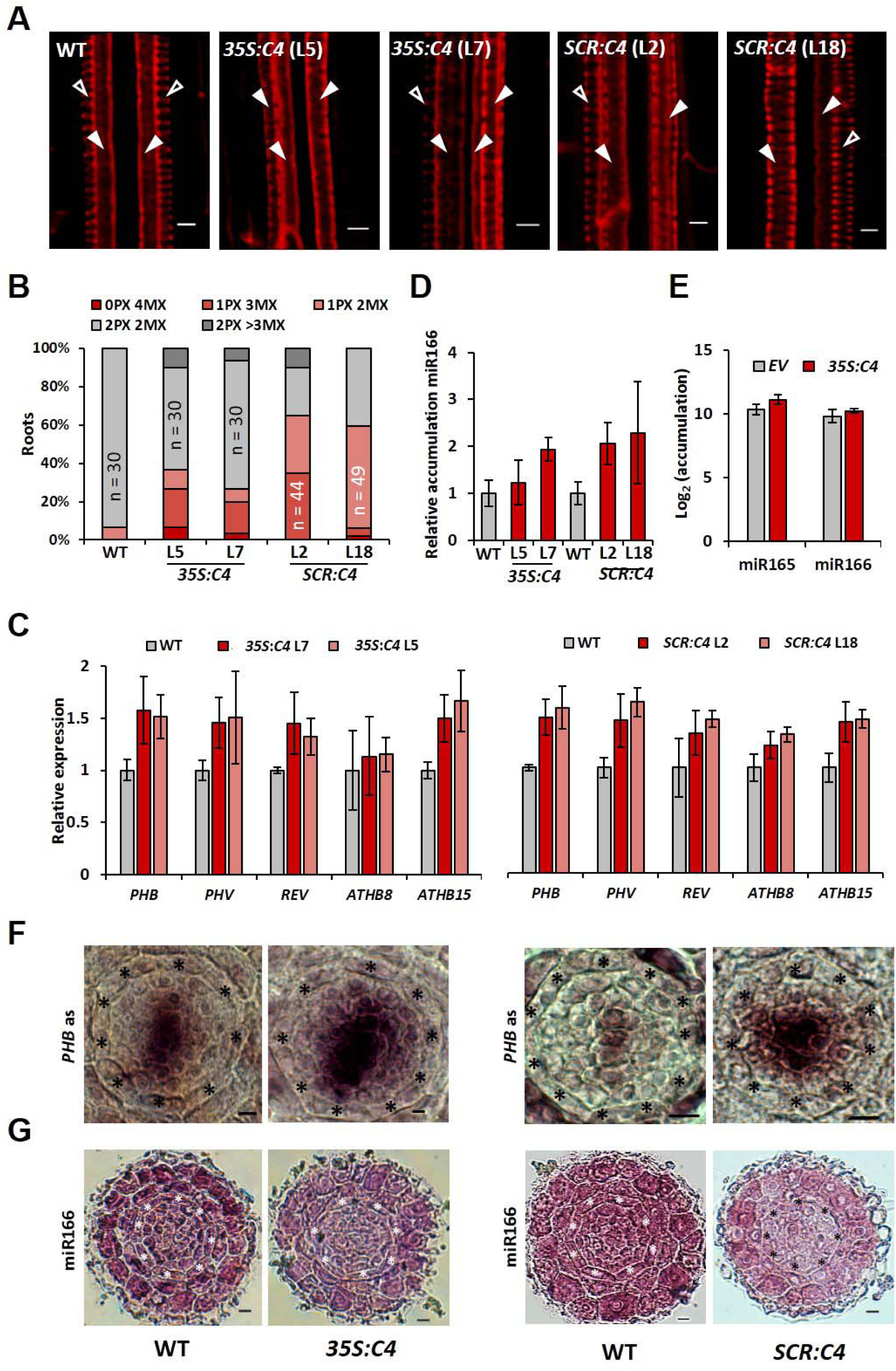
Ubiquitous or endodermis-specific expression of the viral BAM1/2 interactor C4 interferes with the cell-to-cell movement of miR165/6 and xylem development. **A.** Confocal micrographs of basic fuchsin-stained xylem in WT, 35S:*C4*, and *SCR:C4* five-day-old roots. Empty arrowheads indicate protoxylem; filled arrowheads indicate metaxylem. Scale bar = 4µm. **B.** Quantification of the number of protoxylem (PX) and metaxylem files (MX) in the roots of listed genotypes. **C**, **D.** Accumulation of transcripts of the *HD-ZIPIII* family genes (**C**), and miR166 (**E**) in eleven-day-old WT or *35S:C4* seedlings and in five-day-old WT or *SCR:C4* seedlings as measured by qRT-PCR. Results are the average of three biological replicates. Error bars indicate SD. **E.** miR165/6 accumulation in *S-S*/*35S:C4* and *S-S/EV* control as measured by sRNA-seq. **F.** *In situ* hybridization with a *PHB* mRNA specific probe on cross-sections of WT, *35S:C4*, and *SCR:C4* roots. Five *35S:C4* roots and eight *SCR:C4* roots were checked, together with the same number of WT roots; all roots showed a similar phenotype. One representative picture is shown. **G.** *In situ* hybridization with a miRNA166-specific LNA probe on cross-sections of WT, *35S:C4*, and *SCR:C4* roots. Thirteen 35S*:C4* roots and seventeen *SCR:C4* roots were checked; five 35S*:C4* roots and seven *SCR:C4* roots showed the miR166 distribution pattern displayed in this figure, with weaker signal in the stele. Scale bar = 6µm. Asterisks indicate the position of endodermis.

Recently, BAM1 was shown to act as a receptor for the CLE9/10 peptides to regulate periclinal cell division of xylem precursor cells ^11^. The results presented here unveil an additional, novel, redundant role of BAM1 and BAM2 in the regulation of xylem cell fate in the root stele. *bam1 bam2* double mutants display defects in xylem patterning, which are mimicked cell-autonomously and non-cell-autonomously by the expression of the viral BAM1/2-interactor C4; however, all regulatory steps occurring upstream of the cell-to-cell movement of miR165/6 are unaltered in the absence of BAM1/2 or in the presence of C4. Despite normal accumulation of miR165/6, the action of these miRNAs on their target *PHB* is compromised in *bam1 bam2* mutant or C4 transgenic lines, which correlates with a reduced distribution of miR166 in the root stele, underpinning the observed defective xylem patterning. Therefore, BAM1 and BAM2 seem to promote the cell-to-cell movement of both siRNA ^9^ and miRNA, an activity targeted by the viral effector C4; whether their role in sRNA-mediated intercellular communication underlies other biological functions of BAM1/2 remains to be determined.

Although our results provide novel insight into the mechanisms enabling the cell-to-cell movement of sRNA, which virtually impacts every aspect of plant biology, our current view of this process is still extremely limited and multiple questions remain to be answered. For example, whether sRNA travel in a free form or associated to proteins, or how directionality of the movement, if required, is accomplished, are long-standing questions. The elucidation of how BAM1/2 exert their role on the intercellular spread of sRNA at the molecular and cellular levels may shed light on these and other still elusive matters. However, it must be kept in mind that BAM1/2 are likely not the only proteins mediating the cell-to-cell movement of sRNA in plants: considering the restricted expression pattern of *BAM1/2*, together with the limited developmental phenotypes of the *bam1 bam2* double mutants, additional molecular mechanisms must regulate this process outside the *BAM1/2* expression domains.

## METHODS

### Plant materials and growth conditions

Mutants and transgenic plants used in this study are summarized in Table S1. Seedlings used for quantitative RT-PCR (qRT-PCR) and xylem phenotype analysis were grown on half strength Murashige and Skoog (1/2MS) medium containing 1% sucrose and 1% agar. Plates were placed vertically in a growth chamber with a photoperiod of 16 h light/8 h dark at 22°C. *SCR:C4* plants used for phenotyping were grown in soil under the same environmental conditions described above.

### Real-time quantitative RT-PCR (qRT-PCR)

For real-time quantitative RT-PCR (qRT-PCR), total RNA was extracted using Plant RNA Kit (Omega, USA) and reverse-transcribed by First Chain cDNA Synthesis Kit (TonkBio, China). qPCR was performed using C1000 Touch Thermal Cycler (Bio-Rad, USA); 20μl of PCR reaction mixture contained 10μl of SYBR Green mix (Bio-Rad, USA), 1μl of primer mix (10μM), 1μl reverse-transcribed product and 8μl of water. *ACTIN* (*ACT2*) was used as normalizer. Data were analyzed using the 2^ΔΔCT^ method. To quantify the accumulation of miR166, stem-loop qPCR was conducted as previously described^12^. All primers used for qPCR are listed in Table S2.

### Constructs and generation of transgenic lines

To generate the pSCR:*C4* construct, the coding sequence of C4 was cloned into pENTR/D-TOPO (Invitrogen, USA), and subsequently Gateway-cloned into the pSCR:GW vector^13^ through an LR reaction (Invitrogen, USA). *A*. *thaliana* plants were transformed using the floral dipping method^14^.

### In situ hybridization

In situ hybridization was performed as previously described^15,16^. The probe for *PHB* detection was cloned into the pGEM-T Easy vector (Promega, USA), using the primers listed in Table S2. For microRNA in situ hybridization, a specific miR166 LNA probe (QIAGEN, Germany) was used. 100 ng probe were used per slide. The hybridization temperature was 52°C for PHB detection, and 58°C for miR166 detection.

### Small RNA (sRNA) sequencing

Small-RNA (sRNA) data analyses were performed using a pipeline previously described^17^. Briefly, raw reads were trimmed using trim_galore v0.4.0 (https://www.bioinformatics.babraham.ac.uk/projects/trim_galore/) to remove the adapter sequences and bases that have a quality score lower than 10. Reads that could not be aligned to structural RNA sequences (rRNA, tRNA, snoRNA, snRNA, etc.) were aligned to the TAIR10 genome using Burrows–Wheeler aligner by allowing one mismatch per read^17^. The Tair10 genome was divided into non-overlapping 200-bp bins. The number of sRNA reads (with different lengths) in each 200-bp bin or specific genes were summarized and normalized to the structural RNA-removed library size (reads per 10 million) using bedtools v2.26.0 (https://bedtools.readthedocs.io/en/latest/). Results from two independent transgenic lines per construct were pooled.

### Confocal imaging

All confocal images were acquired using a Leica TCS SP8 point scanning confocal microscope. For basic fuchsin staining, 5- or 6-day-old seedlings were first treated with 1M KOH solution for 6 hours at 37°C. Seedlings were then stained with 0.01% basic fuchsin solution in water for 5 minutes, and subsequently destained in 70% ethanol for 10 minutes. To check *BAM1* expression pattern and SHR-GFP movement in the root tip, 5-day-old seedlings were imaged after propidium iodide (PI) staining. The settings used for the laser scanning are as follows: Ex:561nm, Em:600-700nm for basic fuchsin staining; Ex:488nm, Em:500-550nm for GFP; Ex:514nm, Em:525-570nm for YFP; Ex:561nm, Em:-680 nm for PI staining.

## Supporting information

Supplementary references

## ACKNOWLEDGEMENTS

The authors thank Steven Clark and Zachary Nimchuk for kindly sharing materials; Wenjie Zeng, Xinyu Jian, Aurora Luque, and Yujing (Ada) Liu for technical assistance; and all members in Rosa Lozano-Duran’s and Alberto Macho’s groups for stimulating discussions and helpful suggestions. This research was supported by the Strategic Priority Research Program of the Chinese Academy of Sciences, Grant No. XDB27040206, and by the National Natural Science Foundation of China (NSFC) (grant numbers 31671994 and 31870250). Research in RL-D’s lab is funded by the Shanghai Center for Plant Stress Biology of the Chinese Academy of Sciences and the 100 Talent program of the Chinese Academy of Sciences. We apologize to authors of relevant primary research works that could not be directly cited in this manuscript due to length restrictions.

## SUPPLEMENTARY MATERIAL

**Supplementary figure 1. Expression pattern of *BAM1* in the root. A**, **B**. Propidium iodide-stained root of a six-day-old transgenic *pBAM1*:*YFP-NLS* Arabidopsis seedlings. Scale bar = 10 µm (A), 20 µm (B). Asterisks indicate the position of the endodermis. Arrowheads indicate xylem cell files. **C.** Tissue-specific expression of *BAM1* and *BAM2* in roots (images taken from the Arabidopsis eFP browser).

**Supplementary figure 2. *bam1 bam2* double mutants display short roots. A**, **B.** Six-day-old seedlings of *bam1-3 bam2-3* double mutants (**A**) or *SUC:SUL/bam1 bam2* (lines 1.8 and 1.41) (**B**) and their respective controls. WT: wild type (L*er*); S-S: *SUC:SUL*; EV: empty vector. Scale bar = 0.5cm.

**Supplementary figure 3. *bam1* and *bam2* single mutants have normal xylem. A-D.** Basic fuchsin-stained xylem of six-day-old Col-0 WT (**A**), *bam1-3* (**B**), L*er* WT (**C**) and *bam2-3* (**D**). Scale bar = 4µm.

**Supplementary figure 4. Expression of *SCR* and *SHR* is not reduced in the *bam1 bam2* double mutant.** Accumulation of *SHR* and *SCR* transcripts in six-day-old *bam1-3 bam2-3* double mutant roots compared to the WT (L*er*) control, as measured by qRT-PCR. Results are the mean of three biological replicates; error bars indicate SD.

**Supplementary figure 5. Overexpression of *BAM1* has no effect on xylem development.** Accumulation of *BAM1* and *BAM2* (left) and *HD-ZIPIII* family genes (right) transcripts in roots of WT (Col-0) and *35S*:*BAM1-GFP* eleven-day-old seedlings, as measured by qRT-PCR. Results are the mean of three biological replicates; error bars represent SD. **B.** Basic fuchsin-stained xylem of WT (Col-0) and *35S*:*BAM1-GFP* six-day-old roots. Scale bar = 4µm.

**Supplementary figure 6. C4_G2A_ has no effect on xylem development. A.** Accumulation of *HD-ZIPIII* family genes transcripts in WT (Col-0) and *35S*:*C4*_*G2A*_ eleven-day-old seedlings as measured by qRT-PCR. Results are the mean of three biological replicates; error bars represent SD. This experiment was performed together with that shown in Fig. 2C and shares the same control. **B.** Quantification of the number of protoxylem (PX) and metaxylem files (MX) in the roots of 5-day-old *35S*:*C4*_*G2A*_ seedlings. This experiment was performed together with that shown in Fig. 2B and shares the same control.

**Supplementary figure 7. Expression of *SCR* and *SHR* is not reduced in transgenic plants expressing C4. A**, **B.** Accumulation of *SHR* and *SCR* transcripts in roots of eleven-day-old *35S*:*C4* and *35S*:*C4*_*G2A*_ seedlings (**A**), or five-day-old *SCR:C4* seedlings (**B**) compared to the WT (Col-0) control as measured by qRT-PCR. Results are the mean of three biological replicates; error bars indicate SD. **C**, **D.** Accumulation of *MIR166A/B* transcripts in eleven-day-old *35S*:*C4* and *35S*:*C4*_*G2A*_ roots (**C**), or in five-day-old *SCR:C4* seedlings (**D**) compared to the WT (Col-0) control as measured by qRT-PCR. Results are the mean of three biological replicates; error bars indicate SD.

**Supplementary figure 8. C4 does not affect SHR movement. A-C.** Localization of SHR-GFP in transgenic *SHR:SHR-GFP* five-day-old roots in the absence (WT) (**A**) or presence of *SCR:C4* (lines 2 and 18) (**B**, **C**). Scale bar = 20µm. Asterisks indicate the position of the endodermis.

**Supplementary figure 9. Developmental phenotypes of *SCR:C4* plants. A.** Flowering six-week-old plants grown in long day conditions. **B.** Rosettes of four-week-old plants grown in long day conditions. **C.** Representative flowers and siliques. **D.** Typical floral stem. **E-G.** Quantification of stem length (**E**), branch length (**F**), and branch angle (**G**) in WT (Col-0) and *SCR:C4* plants. n=3. Asterisks indicate a statistically significant difference (***, p-value < 0.0001; **, p-value <0.003; *, p-value <0.05), according to a Dunnett test. Scale bar = 2cm.

**Table S1. Plant material used in this study.**

**Table S2. Primers used in this study.**

**Supplementary references.**

