## Supplementary figures for "The receptor-like kinases BAM1 and BAM2 promote the cell-to-cell movement of miRNA in the root stele to regulate xylem patterning"

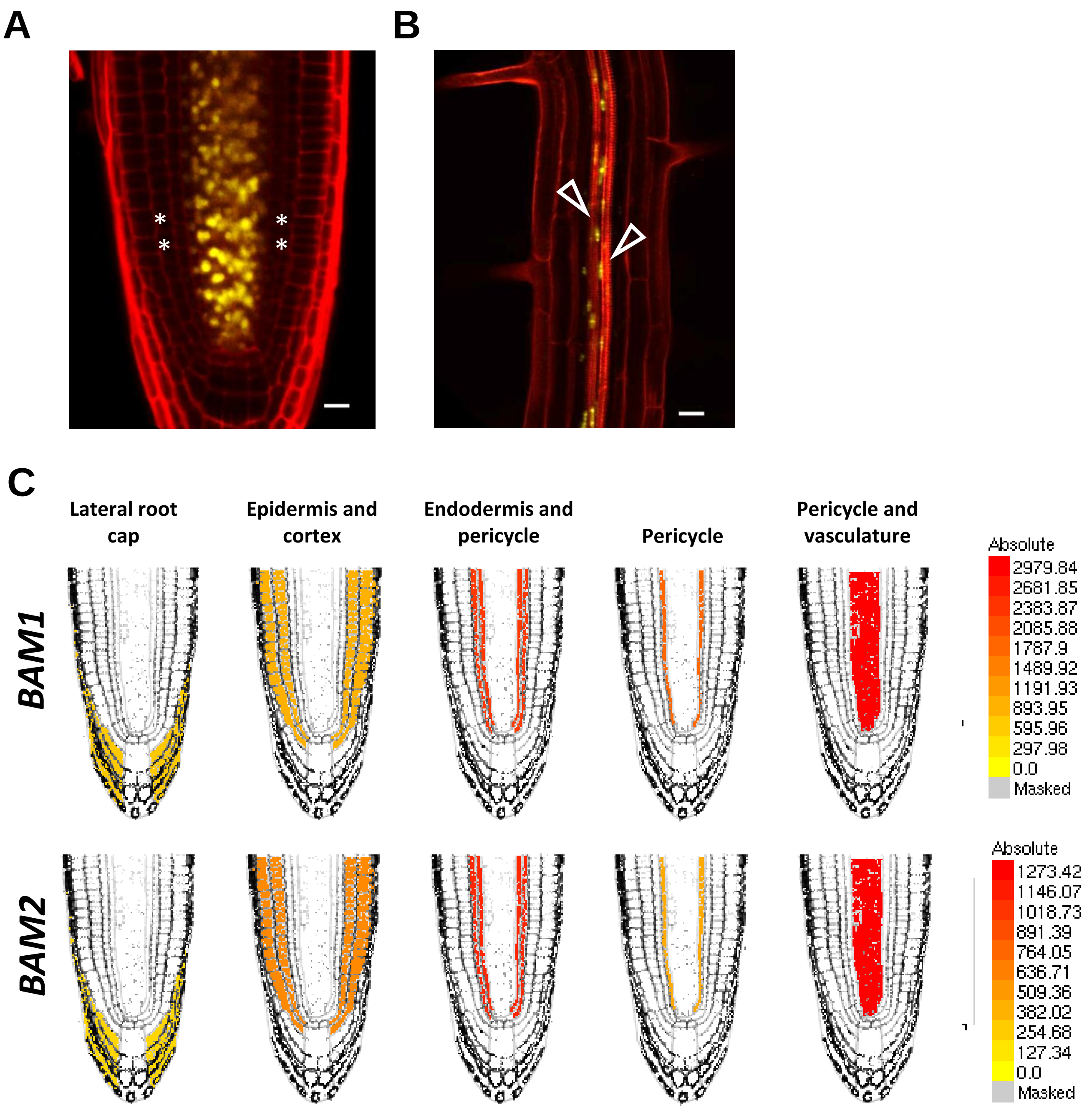


**Supplementary figure 1. Expression pattern of *BAM1* in the root. A, B**. Propidium iodide-stained root of a six-day-old transgenic *pBAM1*:*YFP-NLS* Arabidopsis seedlings. Scale bar = 10 µm (A), 20 µm (B). Asterisks indicate the position of the endodermis. Arrowheads indicate xylem cell files. **C.** Tissue-specific expression of *BAM1* and *BAM2* in roots (images taken from the Arabidopsis eFP browser).

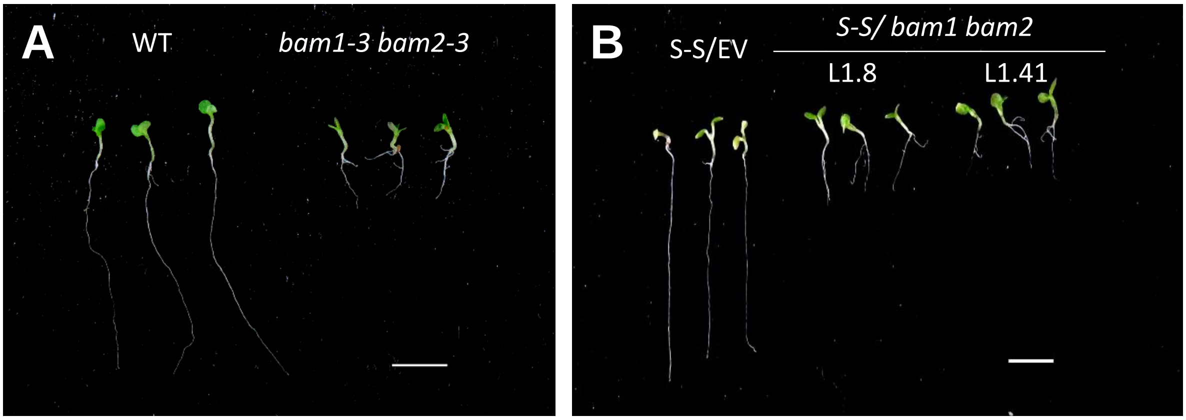


**Supplementary figure 2. *bam1 bam2* double mutants display short roots. A, B.** Six-day-old seedlings of *bam1-3 bam2-3* double mutants (**A**) or *SUC:SUL/bam1 bam2* (lines 1.8 and 1.41) (**B**) and their respective controls. WT: wild type (L*er*); S-S: *SUC:SUL*; EV: empty vector. Scale bar = 0.5cm.

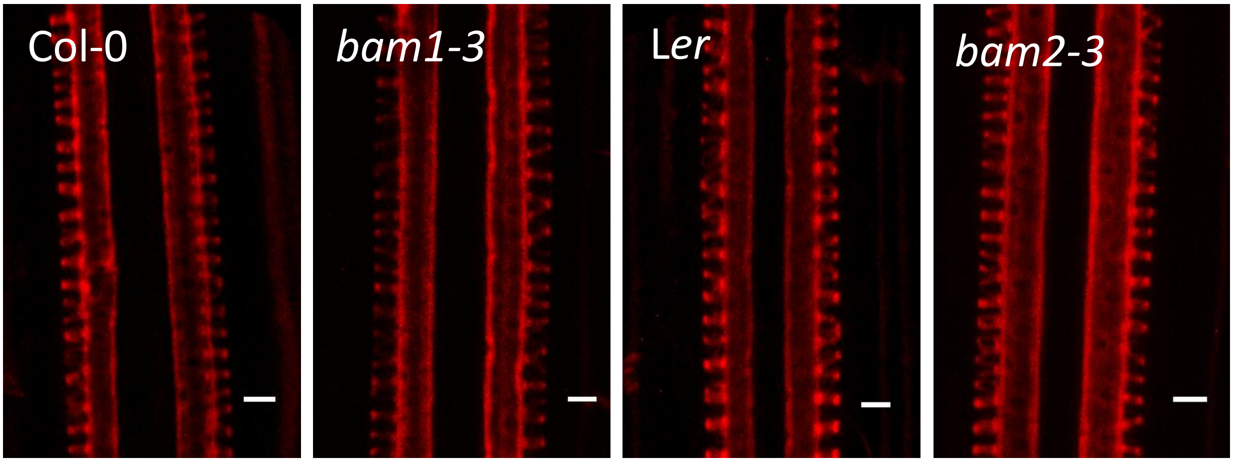


**Supplementary figure 3. *bam1* and *bam2* single mutants have normal xylem. A-D.** Basic fuchsin-stained xylem of six-day-old Col-0 WT (**A**), *bam1-3* (**B**), L*er* WT (**C**) and *bam2-3* (**D**). Scale bar = 4µm.

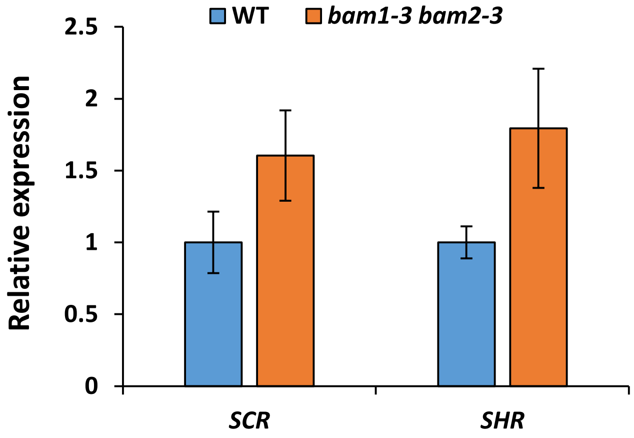


**Supplementary figure 4. Expression of *SCR* and *SHR* is not reduced in the *bam1 bam2* double mutant.** Accumulation of *SHR* and *SCR* transcripts in six-day-old *bam1-3 bam2-3* double mutant roots compared to the WT (L*er*) control, as measured by qRT-PCR. Results are the mean of three biological replicates; error bars indicate SD.

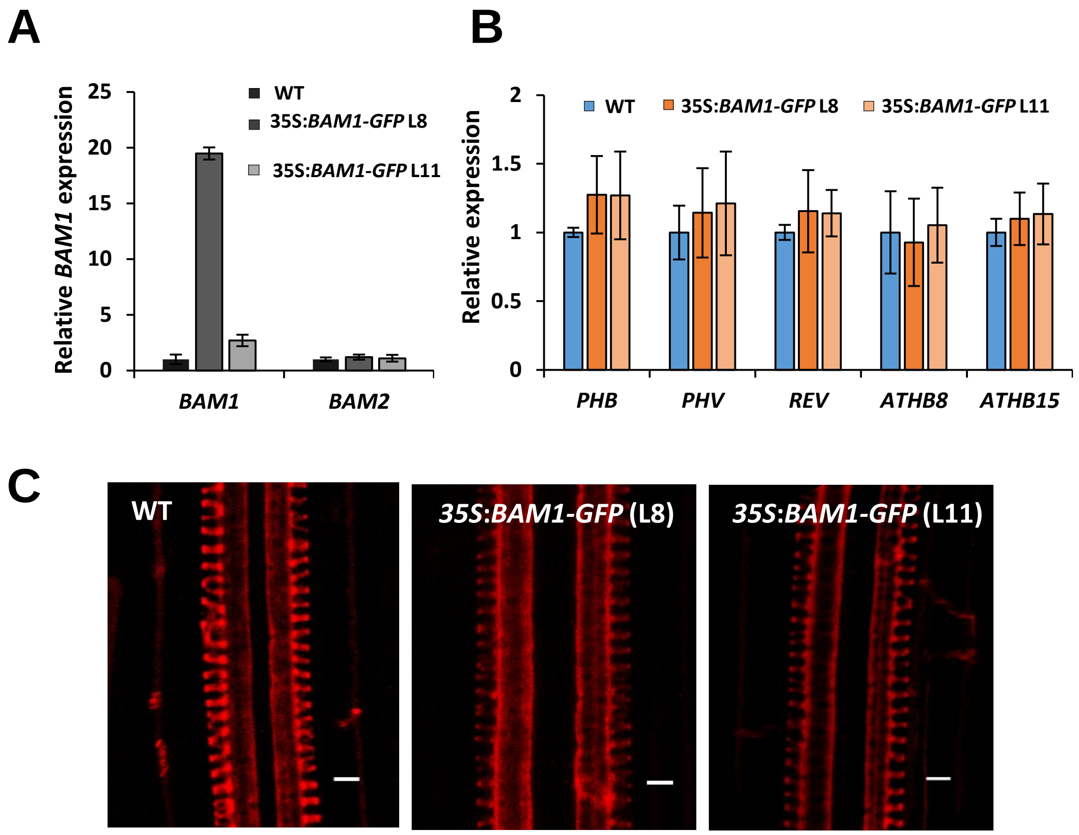


**Supplementary figure 5. Overexpression of *BAM1* has no effect on xylem development. A.** Accumulation of *BAM1* and *BAM2* (left) and *HD-ZIPIII* family genes (right) transcripts in roots of WT (Col-0) and *35S*:*BAM1-GFP* eleven-day-old seedlings, as measured by qRT-PCR. Results are the mean of three biological replicates; error bars represent SD. **B.** Basic fuchsin-stained xylem of WT (Col-0) and *35S*:*BAM1-GFP* six-day-old roots. Scale bar = 4µm.

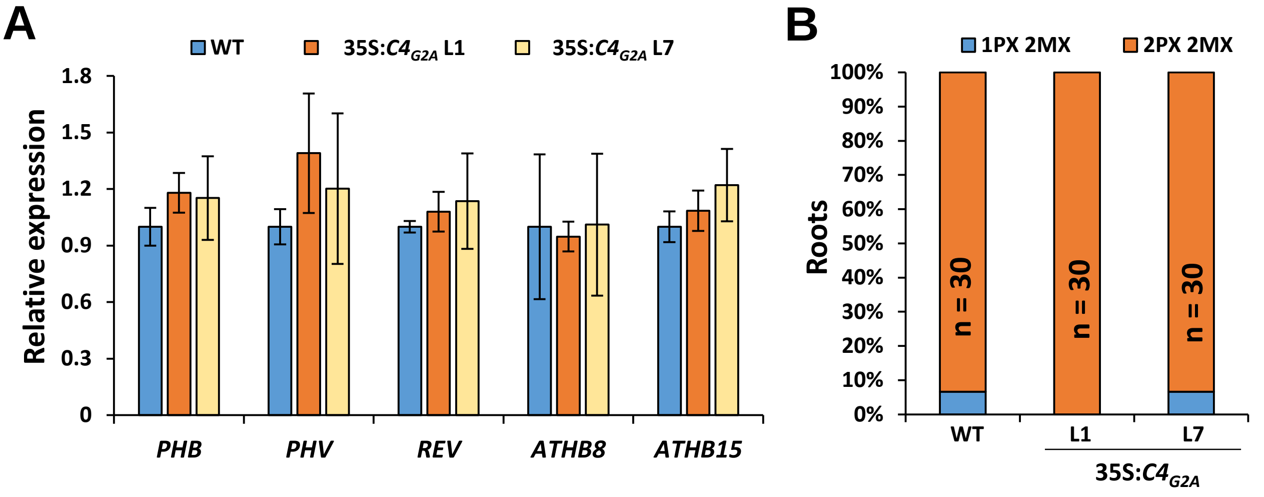


**Supplementary figure 6. C4$_{G2A}$ has no effect on xylem development. A.** Accumulation of *HD-ZIPIII* family genes transcripts in WT (Col-0) and *35S*:*C4*$_{G2A}$ eleven-day-old seedlings as measured by qRT-PCR. Results are the mean of three biological replicates; error bars represent SD. This experiment was performed together with that shown in Fig. 2C and shares the same control. **B.** Quantification of the number of protoxylem (PX) and metaxylem files (MX) in the roots of 5-day-old *35S*:*C4*$_{G2A}$ seedlings. This experiment was performed together with that shown in Fig. 2B and shares the same control.

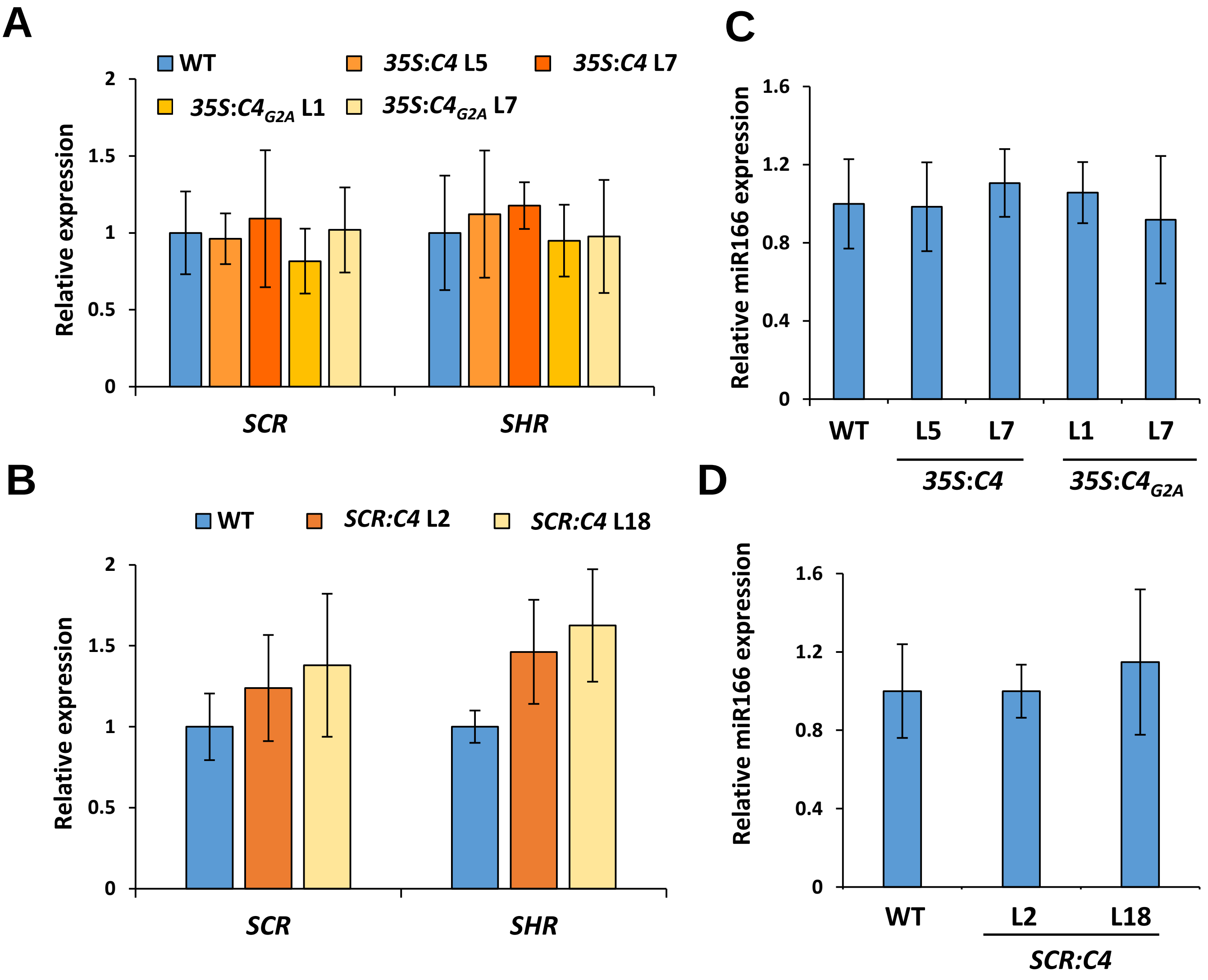


**Supplementary figure 7. Expression of *SCR* and *SHR* is not reduced in transgenic plants expressing C4. A, B.** Accumulation of *SHR* and *SCR* transcripts in roots of eleven-day-old *35S:C4* and *35S:C4*$_{G2A}$ seedlings (**A**), or five-day-old *SCR:C4* seedlings (**B**) compared to the WT (Col-0) control as measured by qRT-PCR. Results are the mean of three biological replicates; error bars indicate SD. **C, D.** Accumulation of *MIR166A/B* transcripts in eleven-day-old *35S:C4* and *35S:C4*$_{G2A}$ roots (**C**), or in five-day-old *SCR:C4* seedlings (**D**) compared to the WT (Col-0) control as measured by qRT-PCR. Results are the mean of three biological replicates; error bars indicate SD.

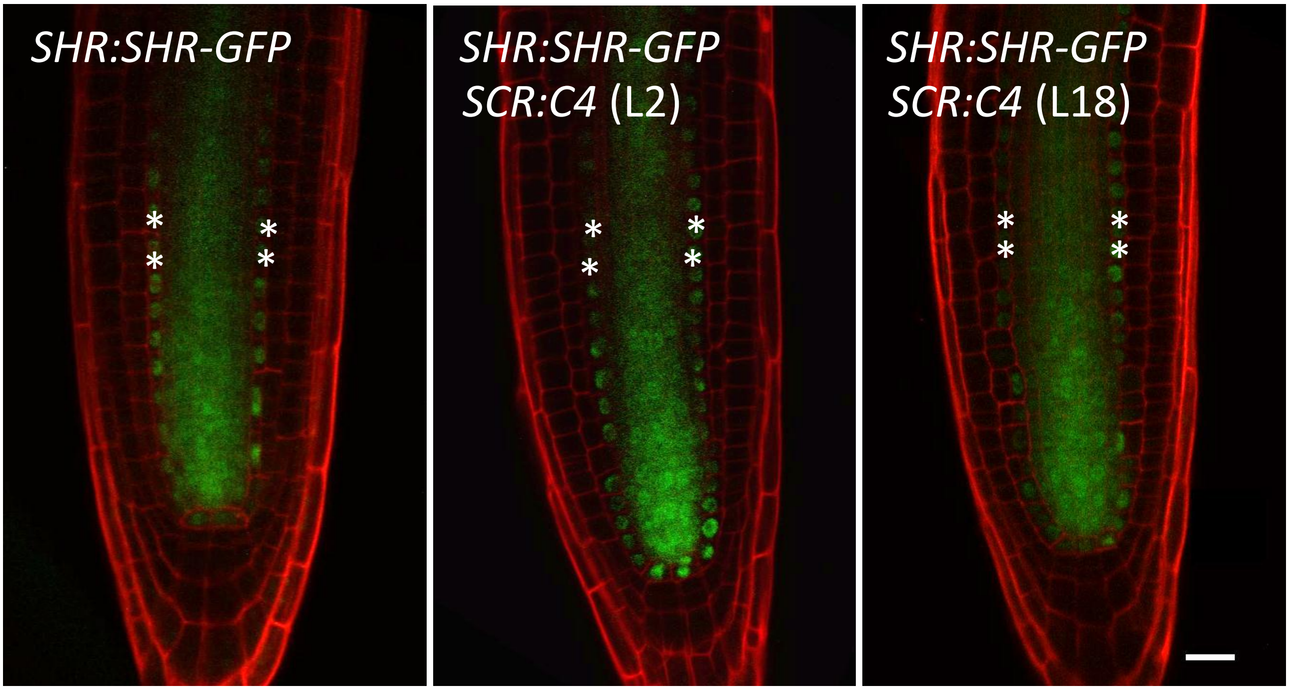


**Supplementary figure 8. C4 does not affect SHR movement. A-C.** Localization of SHR-GFP in transgenic *SHR:SHR-GFP* five-day-old roots in the absence (WT) (**A**) or presence of *SCR:C4* (lines 2 and 18) (**B, C**). Scale bar = 20µm. Asterisks indicate the position of the endodermis.

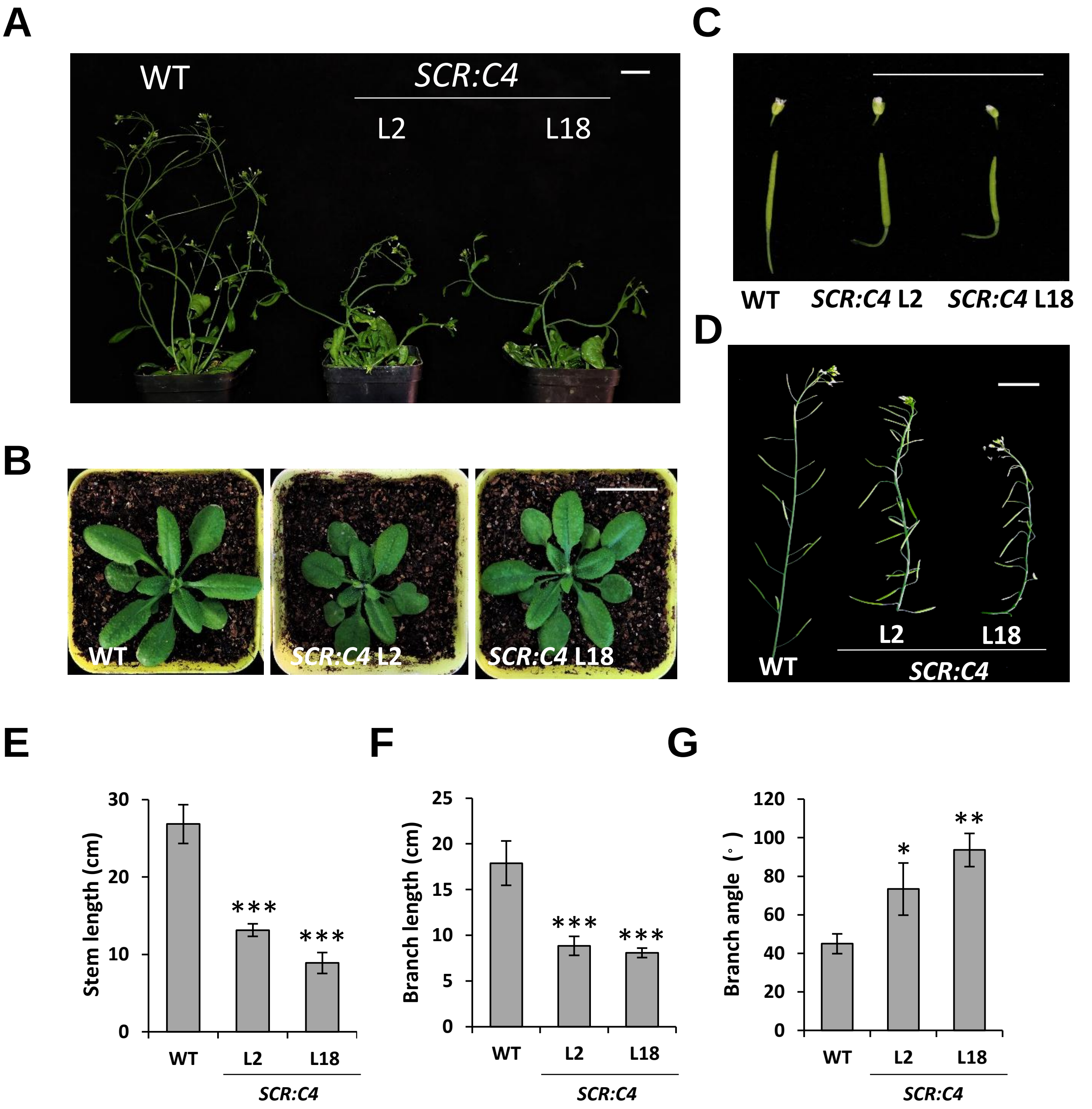


**Supplementary figure 9. Developmental phenotypes of *SCR:C4* plants. A.** Flowering six-week-old plants grown in long day conditions. **B.** Rosettes of four-week-old plants grown in long day conditions. **C.** Representative flowers and siliques. **D.** Typical floral stem. **E-G.** Quantification of stem length (**E**), branch length (**F**), and branch angle (**G**) in WT (Col-0) and *SCR:C4* plants. n=3. Asterisks indicate a statistically significant difference (***, p-value < 0.0001; **, p-value <0.003; *, p-value <0.05), according to a Dunnett test. Scale bar = 2cm.
