## Supplementary material for "The receptor-like kinases BAM1 and BAM2 promote the cell-to-cell movement of miRNA in the root stele to regulate xylem patterning": Table S1

**Table S1. Plant material used in this study.**

| Name | Ecotype | Reference |
| --- | --- | --- |
| *35S:C4* L5 | Col-0 | Rosas-Diaz *et al*., 2018 |
| *35S:C4* L7 | Col-0 | Rosas-Diaz *et al*., 2018 |
| *SCR:C4* L2 | Col-0 | This study |
| *SCR:C4* L18 | Col-0 | This study |
| *bam1-3* | Col-0 | DeYoung *et al*., 2006 |
| *bam2-3* | L*er* | DeYoung *et al*., 2006 |
| *bam1-3 bam2-3* | Introgression of Col-0 into L*er* | DeYoung *et al*., 2006 |
| S-S/CRISPR-CAS9 *bam1 bam2* L1.8 | Col-0 | Rosas-Diaz *et al*., 2018 |
| S-S/CRISPR-CAS9 *bam1 bam2* L1.41 | Col-0 | Rosas-Diaz *et al*., 2018 |
| *pBAM1*:*YFP-NLS* | Col-0 | Rosas-Diaz *et al*., 2018 |
| *pSHR*:*SHR-GFP* | Ws | Nakajima *et al*., 2001 |
