## Supplementary material for "The receptor-like kinases BAM1 and BAM2 promote the cell-to-cell movement of miRNA in the root stele to regulate xylem patterning": Table S2

**Table S2. Primers used in this study.**

| Name | Sequence (5'-3') | Reference | Purpose |
| --- | --- | --- | --- |
| qPHB-F | CTTTGGTAGTGGCGTGCTTT | This study | qPCR for *PHB* |
| qPHB-R | GCCCATTCAGATCGGTGTTC | This study | qPCR for *PHB* |
| qPHV-F | CCAAGATCATGCAGCAGGGA | This study | qPCR for *PHV* |
| qPHV-R | CGCTTGCTCATACGAAACCG | This study | qPCR for *PHV* |
| qREV-F | AGAGCATCGATCTGAGTGGG | This study | qPCR for *REV* |
| qREV-R | TTGTTGGTCTCATTCCCGGA | This study | qPCR for *REV* |
| qATHB15-F | AGAATGTTCCTCCGGCGATC | This study | qPCR for *ATHB15* |
| qATHB15-R | TGCCCTCCAAATCCTCCAAC | This study | qPCR for *ATHB15* |
| qATHB8-F | TGTTGCTCACTCAAGGCCTT | This study | qPCR for *ATHB8* |
| qATHB8-R | TCTTGAAGTGCCACCAACGT | This study | qPCR for *ATHB8* |
| ACT2-F | CTAAGCTCTCAAGATCAAAGGCTTA | McKinney and Meagher, 1998 | qPCR for *ACTIN* |
| ACT2-R | ACTAAAACGCAAAACGAAAGCGGTT | McKinney and Meagher, 1998 | qPCR for *ACTIN* |
| qPri-MIR166-F | TGTCTGGCTCGAGGACTCTT | This study | qPCR for pri-miR166a |
| qPri-MIR166-R | TCCGACGACACTAAAACCCT | This study | qPCR for pri-miR166a |
| RT-primer | GTCGTATCCAGTGCAGGGTCCGAGGTATTCGCACTGGATACGACGGGGAA | Varkonyigasic *et al.*, 2007 | Stem loop qPCR for miR166 |
| qmiR166-F | TCGCTTCGGACCAGGCTTCA | Varkonyigasic *et al.*, 2007 | Stem loop qPCR for miR166 |
| qmiR166-R | GTGCAGGGTCCGAGGT | Varkonyigasic *et al.*, 2007 | Stem loop qPCR for miR166 |
| qSHR-F | GGCGGATGATGTCAGAGCTT | This study | qPCR for *SHR* |
| qSHR-R | CAAACCACCGGCTGATCTCT | This study | qPCR for *SHR* |
| qSCR-F | CAGCAGCACCAACAACAACA | This study | qPCR for *SCR* |
| qSCR-R | GGTGGTGCATCGGTAGAAGA | This study | qPCR for *SCR* |
| qBAM1-F | TCCAATAAGCTGACCGGAAC | This study | qPCR for *BAM1* |
| qBAM1-R | CCGGGTCAAAGACTCACATT | This study | qPCR for *BAM1* |
| qBAM2-F | ATGTTTACCGGCGAGATTCC | This study | qPCR for *BAM2* |
| qBAM2-R | GCACCGTAGAGCTTATTCCTGA | This study | qPCR for *BAM2* |
| pSCR：GW-F | GGAGATCGTGAAGACGATCAAG | This study | Genotyping p*SCR:C4* trangenic lines |
| C4-S-R | TTAATATATTGAGGGCCTCGG | This study | Genotyping p*SCR:C5* trangenic lines |
| BAM2 5´3 | CAATTACCTTACCGGAGAGTTG | DeYoung *et al.*, 2006 | Genotyping *bam2-3* |
| BAM2 3'10 | GGACATTGTAGCCAATCGTTTG | DeYoung *et al.*, 2006 | Genotyping *bam2-3* |
| Ds3-1 | ACCCGACCGGATCGTATCGGT | DeYoung *et al.*, 2006 | Genotyping *bam2-3* |
| BAM1-F | CACCATGAAACTTTTTCTTCTCCT | Rosas-Diaz *et al.*, 2018 | Genotyping *bam1bam2* mutant by CRISPR-CAS9 |
| BAM1-R | GAAAGCGTTGTAGTAGCCGAT | Rosas-Diaz *et al.*, 2018 | Genotyping *bam1bam2* mutant by CRISPR-CAS9 |
| BAM2-F | CACCATGAAGCTTCTTCTTC | Rosas-Diaz *et al.*, 2018 | Genotyping *bam1bam2* mutant by CRISPR-CAS9 |
| BAM2-R | CGTTAGGTTTCCGATCTCCG | Rosas-Diaz *et al.*, 2018 | Genotyping *bam1bam2* mutant by CRISPR-CAS9 |
| PHB-F | ACAGAAATCTACTCCGAACGGTGC | Smith and Long, 2010 | Synthesizing *PHB* probes for in situ hybridization |
| PHB-R | TGCCTGCTCGTAAGATACCATC | Smith and Long, 2010 | Synthesizing *PHB* probes for in situ hybridization |
